# Biological autoluminescence for assessing oxidative processes in yeast cell cultures

**DOI:** 10.1101/2020.11.19.388801

**Authors:** Petra Vahalová, Kateřina Červinková, Michal Cifra

## Abstract

Nowadays, modern medicine is looking for new, more gentle, and more efficient diagnostic methods. A pathological state of an organism is often closely connected with increased amount of reactive oxygen species. They can react with biomolecules and subsequent reactions can lead to very low endogenous light emission (biological autoluminescence -BAL). This phenomenon can be potentially used as a non-invasive and low-operation-cost tool for monitoring oxidative stress during diseases. To contribute to the understanding of the parameters affecting BAL, we analyzed the BAL from yeast *Saccharomyces cerevisiae* as a representative eukaryotic organism. The relationship between the BAL intensity and the amount of reactive oxygen species that originates as a result of the Fenton reaction as well as correlation between spontaneous BAL and selected physical and chemical parameters (pH, oxygen partial pressure, and cell concentration) during cell growth were established. Our results contribute to real-time non-invasive methodologies for monitoring oxidative processes in biomedicine and biotechnology.

## Introduction

One of the main missions of modern society is to prevent, manage, and cure diseases. A key requirement to achieve this is the development of effective and ideally non-invasive diagnostic methods. New gentle diagnostic methods will help to overcome limitations of current methods, which can be painful for the patient and destructive for the analyzed biosamples. In addition to human diseases, the development of new diagnostic methods is also important for monitoring diseases of animals or plants, which have a strong impact on agriculture and the food industry. In this work, we contribute to the development of a non-invasive biophotonic method for real-time monitoring of oxidative processes in cell cultures as model organisms.

Multiple diseases, including those that are the leading causes of deaths worldwide, such as cancer^1^, neurodegenerative^2^, and cardiovascular^3,4^ diseases, are closely related to an increased amount of reactive oxygen species (ROS) in affected body parts. In living organisms, incomplete reduction of O_2_ during respiration is considered to be a typical source of ROS. However, ROS are also naturally produced during fatty acid metabolism in peroxysomes and by NADPH oxidases in the membrane of macrophages^5^. Non-mitochondrial ROS sources, originating from proteins encoded by NADPH oxidase orthologs, have also been recently discovered in single cell organisms, such as yeast^6^. Therefore, organisms have developed various defense mechanisms to eliminate ROS or prevent the damage caused by them. However, an imbalance between ROS production and the capability of an organism’s defense system to eliminate them leads to oxidative stress and harm of an organism. Interactions of ROS with biomolecules lead to a cascade of reactions. One of the reaction branches leads to the formation of unstable high-energy intermediates such as dioxetanes^7^ and tetraoxides^8^, which decompose to generate electronically excited species^9^. Redundant energy can be released in the form of heat or as light emission^10^. The probabilities of reactions leading to the formation of excited species and subsequent radiative de-excitation are low; therefore, the intensity of photon emission is also very low^9^. In addition to biological autoluminescence (BAL), other terms such as ultra-weak photon emission, biophotonics, and bio-chemiluminescence are used for this phenomenon^10^. Thus, the reaction rate of ROS with biomolecules in a studied sample closely correlates with the BAL intensity. Although the probable general biochemical and biophysical mechanisms of BAL are rather well established^9^, the origin and physiology of the processes leading to BAL are different for different organisms in different conditions^11–14^. The amount of ROS in organisms can be affected by endogenous processes but also by the presence of external oxidants and pro-oxidants. Natural and artificial processes can lead to the generation of various types of ROS. One of the common reactions that yields highly reactive hydroxyl radicals besides other products^15^ is the Fenton reaction, which involves divalent iron (Fe^2+^) and hydrogen peroxide (H_2_O_2_). In organisms, these two chemicals are strictly controlled and separated. However, artificial usage of the Fenton reaction is popular in many industrial fields in which production of the highly reactive radical is desirable.

As a biological system for the analysis of ROS-BAL correlations, we chose yeast *Saccharomyces cerevisiae*, which is a widely used and well characterized model organism in molecular biology research. Furthermore, yeasts and their various strains are also essential in the food industry^16,17^ and biotechnology^18^. They are eukaryotic, which is the same as humans, animals, or plants, but single-celled organisms; thus, it is easier to evaluate the effects of various stress factors on the organism.

There have been several studies of spontaneous BAL from yeast. For example, Quickenden et al.^11,19–22^ studied the intensity and spectral distribution of spontaneous BAL in the exponential and stationary phase of yeast growth. Other researchers have used various methods to study oxidation caused by the Fenton reaction^23^,24. Ivanova et al.^25^ focused on mechanisms of chemiluminescence as a result of the Fenton reaction. To connect these topics and broaden the knowledge about them, we decided to evaluate the correlation between BAL of a living organism, yeast *Saccharomyces cerevisiae*, and artificially increased amount of ROS by the Fenton reaction. We also investigated the correlation between spontaneous BAL and selected physical and chemical parameters (pH, oxygen partial pressure, cell concentration, and concentration of the antioxidant ascorbic acid) to verify the hypothesis of ROS-BAL correlation.

## Results and Discussion

### Biological autoluminescence enhanced by Fenton reagents

First, we evaluated the BAL signal from yeast samples containing three different cell concentrations to determine which concentration resulted in the highest BAL intensity. The BAL intensity was observed during real-time oxidation of yeast cell samples with 0.5 mM FeSO_4_ and 2.5 mM H_2_O_2_ (final concentrations are always provided, unless noted otherwise). The kinetics during the first 60 s are shown in Figure 2a. Hydrogen peroxide was injected into the sample after 10 seconds of background measurement. The sums of the BAL signal during the first 60 s after injection of H_2_O_2_ with a subtracted background (first 10 s before the injection of H_2_O_2_) are shown in Figure 2b. The average from 3 independent measurements and standard deviation are displayed. We can see that the BAL signal increases together with the yeast concentration up to the limit 10^8^ cells· mL^−1^. With increasing yeast concentration, the amount of biomass that ROS can react with is higher and the amount of potential sources of BAL grows. However, above 10^8^ cells ·mL^−1^, the sample starts to become increasingly turbid and the samples also sediment faster, making the lower layers inaccessible for oxygen and the Fenton reagents, hence decreasing the actual amount of biomass that could be directly oxidised. All these factors, we believe, lead to the lower BAL intensity for cell concentrations above 10^8^ cells· mL^−1^. Therefore, for subsequent experiments we used a yeast concentration of 10^8^ cells· mL^−1^, which exhibited the highest BAL intensity.

**Figure 1.**
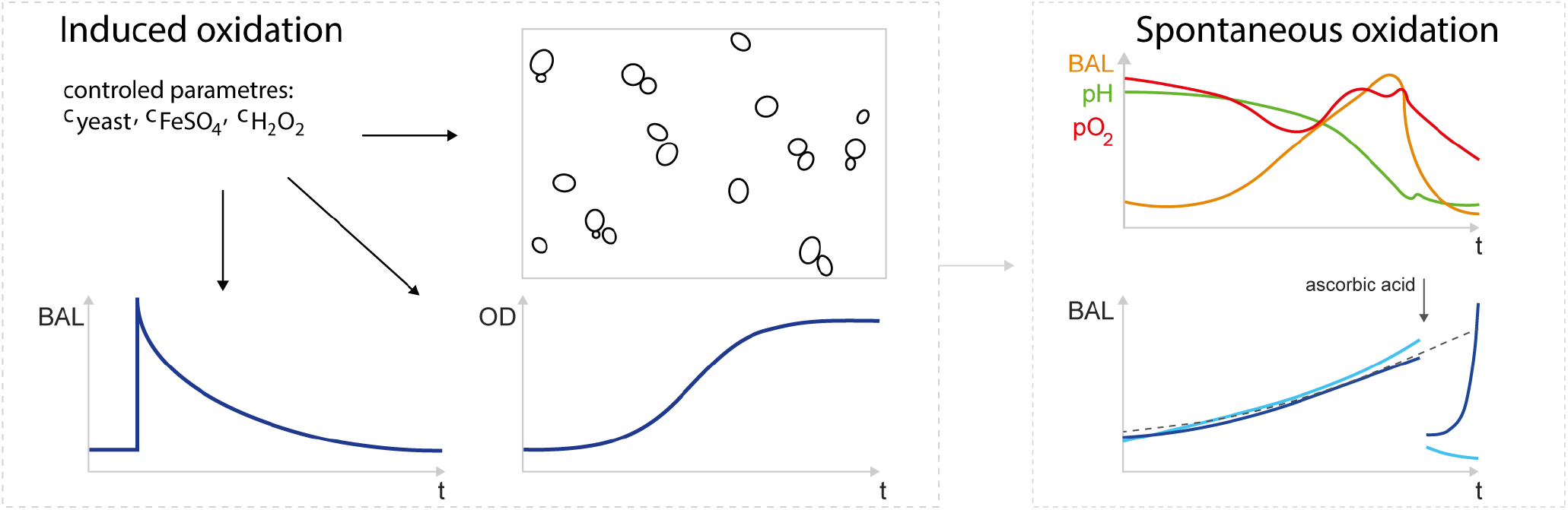
A schematic summary of the parameters and processes related to biological autoluminescence (BAL) explored in this paper. We analyzed the BAL and related parameters both under the conditions of induced and spontaneous (endogenous) oxidation in the yeast cell cultures. For induced oxidation, the amount of free radicals in a sample was regulated (increased) by the Fenton reaction. Then, the effects of various concentration of yeast and Fenton reagents (FeSO_4_ and H_2_O_2_) on BAL, growth curves, and cell clustering was analyzed. The BAL due to spontaneous oxidation processes and its correlation with several selected parameters (pH, pO_2_, concentration of antioxidant ascorbic acid) were established.

**Figure 2.**
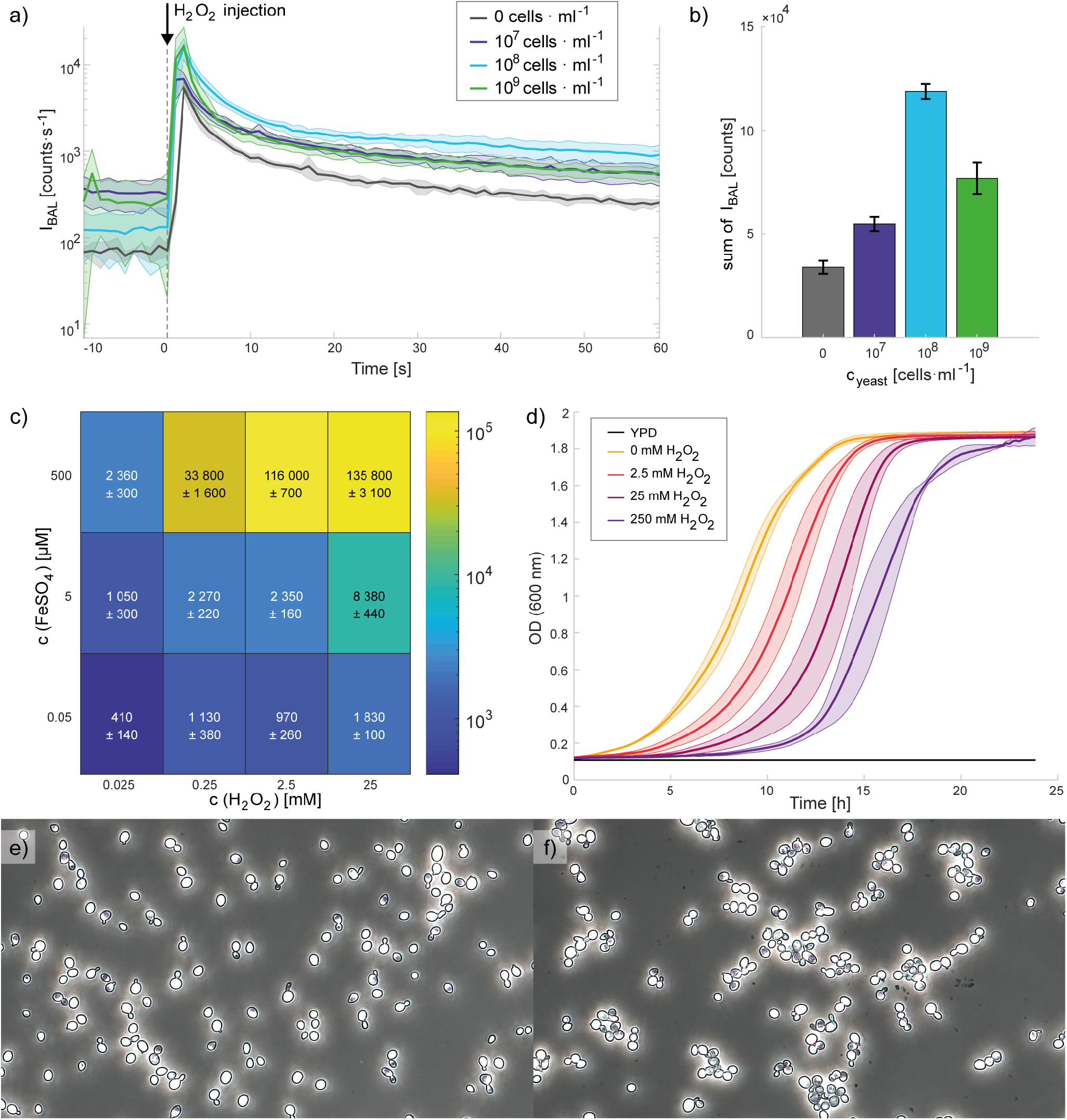
Effects of the Fenton reaction on BAL of yeast, their growth curve, and evaluation of the oxidised yeast sample under a microscope. First two graphs (**a, b**) show induced BAL of yeast by the Fenton reaction (0.5 mM FeSO_4_, 2.5 mM H_2_O_2_) for different concentrations of yeast (10^7^ - 10^9^ cells· mL^−1^). Part **c)** displays induced yeast BAL by the Fenton reaction for various concentrations of Fenton reagents (50 nM – 0.5 mM FeSO_4_, 25 *µ*M – 25 mM H_2_O_2_) and a fixed yeast concentration (10^8^ cells· mL^−1^). **a)** BAL kinetics after the injection of hydrogen peroxide into the sample after 10 seconds of background measurement (t = 0). **b)** Bar graph and **c)** heat map of the sum of the BAL intensities during the first 60 s after injection of H_2_O_2_ with a subtracted background (the previous 10 s before the injection of H_2_O_2_). **d)** Yeast growth curves after oxidation of yeast samples (10^8^ cells mL^−1^) by the Fenton reaction (∼10 min; 0.5 mM FeSO_4_, 0 / 2.5 / 25 / 250 mM H_2_O_2_) depicted by OD at 600 nm. In all cases (a - d), the average from 3 measurements and the standard deviation are shown. Microscopic pictures of a yeast sample in water **e)** before and **f)** after the Fenton reaction (∼10 min; 0.5 mM FeSO_4_, 2.5 mM H_2_O_2_). Yeast concentrations (3 – 9 *µ*m in a diameter) were 10^8^ cells·mL^−1^ before and 3·10^7^ cells·mL^−1^ after oxidation.

We measured the BAL response of yeast samples at a concentration of 10^8^ cells· mL^−1^ induced by the addition of various concentrations of the Fenton reagents FeSO_4_ and H_2_O_2_. The final concentration ranges of the reagents were 50 nM – 0.5 mM FeSO_4_ and 25 *µ*M – 25 mM H_2_O_2_. The sums of the BAL intensities during the first 60 s after injection of H_2_O_2_ with a subtracted background (calculated from the previous 10 s before the injection of H_2_O_2_) are show in Figure 2c. The given numbers (represented by colours) are the average from 3 measurements with a standard deviation. At the lowest used concentrations (50 nM FeSO_4_ and 25 *µ*M H_2_O_2_), a signal comparable with the intensity of a control sample (only milli-Q water without yeast) was obtained, see Fig S1. For the measurements at higher concentrations, the BAL signals were substantially higher than those obtained with the control samples. Generally, higher Fenton reagent concentrations resulted in a higher BAL signal.

The influence of oxidation induced by the Fenton reaction (caused by the highest previously used concentrations 0.5 mM FeSO_4_, 2.5 mM and 25 mM H_2_O_2_) on a yeast sample was evaluated under a microscope. In Fig. 2e,f yeast e) before and f) after 10 min oxidation with 0.5 mM FeSO_4_, 2.5 mM H_2_O_2_ are shown. The yeast concentration (evaluated for cells with a diameter between 3 – 9 *µ*m) was 10^8^ cells· mL^−1^ for control sample and before oxidation (Fig. 2e) and 3 10^7^ cells· mL^−1^ after oxidation of the sample (Fig. 2f). For both H_2_O_2_ concentrations, we did not observe any damaged cells. However, the yeast cells tended to cluster after oxidation. Cell aggregation could be one of the possible reasons for the lower yeast concentration that was measured in the 3 – 9 *µ*m cell diameter range.

Toledano et al. claim that the toxic concentration of hydrogen peroxide for yeast is approximately 5 mM (depending on the yeast species) [5, chap. 6, p.246], which is within the range of the concentrations we evaluated. However, inducing the apoptosis by external oxidants depends on the incubation time. There are also several factors that can modulate the resistance to oxidative stress. For example, yeasts have a higher resistance to oxidative stress in the stationary phase and during respiration, when their metabolic processes (oxidative phosphorylation) employ oxygen and inevitably generate ROS endogenously. Also, previous mild stress condition (including different types of stress) can lead to increased tolerance to oxidative stress. For example, the sudden transfer of yeast cells from room temperature to 30 ^°^C and change of the medium can be experienced as stress [5, chap. 6]. Exploring these factors on oxidation and BAL response might be a part of a future work.

The effect of the Fenton reaction (≈10 min incubation; 0.5 mM FeSO_4_, 2.5 - 250 mM H_2_O_2_) on yeast growth were also evaluated. As shown in Fig. 2d, the samples treated with a higher concentration of hydrogen peroxide had a longer lag phase. However, the final amount of cells (in the stationary phase) was similar for all samples. The duration of the lag phase depends on several factors such as the temperature, aeration, the conditions in a previous environment, amount of transferred biomass, or yeast health [26, chap. 9]. More stressed cells probably need more time to overcome unfavourable consequences and to prepare for cell division.

### Spontaneous biological autoluminescence

We also analyzed the spontaneous BAL of yeast culture in a bioreactor during their growth (Fig. 3a. Because the sample was prepared under normal laboratory light conditions, we observed an initial slight decrease of the BAL signal while the sample was adapting to the dark condition in a black box. Then, the signal increased during several hours. At the same time, the pH decreased and pO_2_ displayed nonlinear behavior. After approximately 12 hours of measurement, the BAL signal sharply decreased, whereas the pH and pO_2_ slightly increased. Quickenden et al. observed that the BAL of yeast *Saccharomyces cerevisiae* was similar to nutrient luminescence (i.e. close to background) in anaerobic conditions (nitrogen instead of air)^21^. However, there was still enough oxygen in our samples after approximately 12 hours (Fig 3a). As shown in Fig. 2d, a transition from exponential to stationary phase of the cell growth curve occurs after approximately 12 hours (in a control sample = without Fenton reagents treatment). This change of phases is called the diauxic shift and it is connected with a change in yeast metabolism from fermentation to aerobic respiration. Therefore, a sharp decrease of the BAL intensity could be caused by internal changes related with the initiation of respiration. This statement might look contradictory at first sight, because within classical understanding of physiological processes underlying BAL, the higher respiration rate (mitochondria-enabled) should lead to a higher rate of ROS generation^27^–30, hence to a higher BAL intensity. To explain our observations, we propose the following hypothesis:

**Figure 3.**
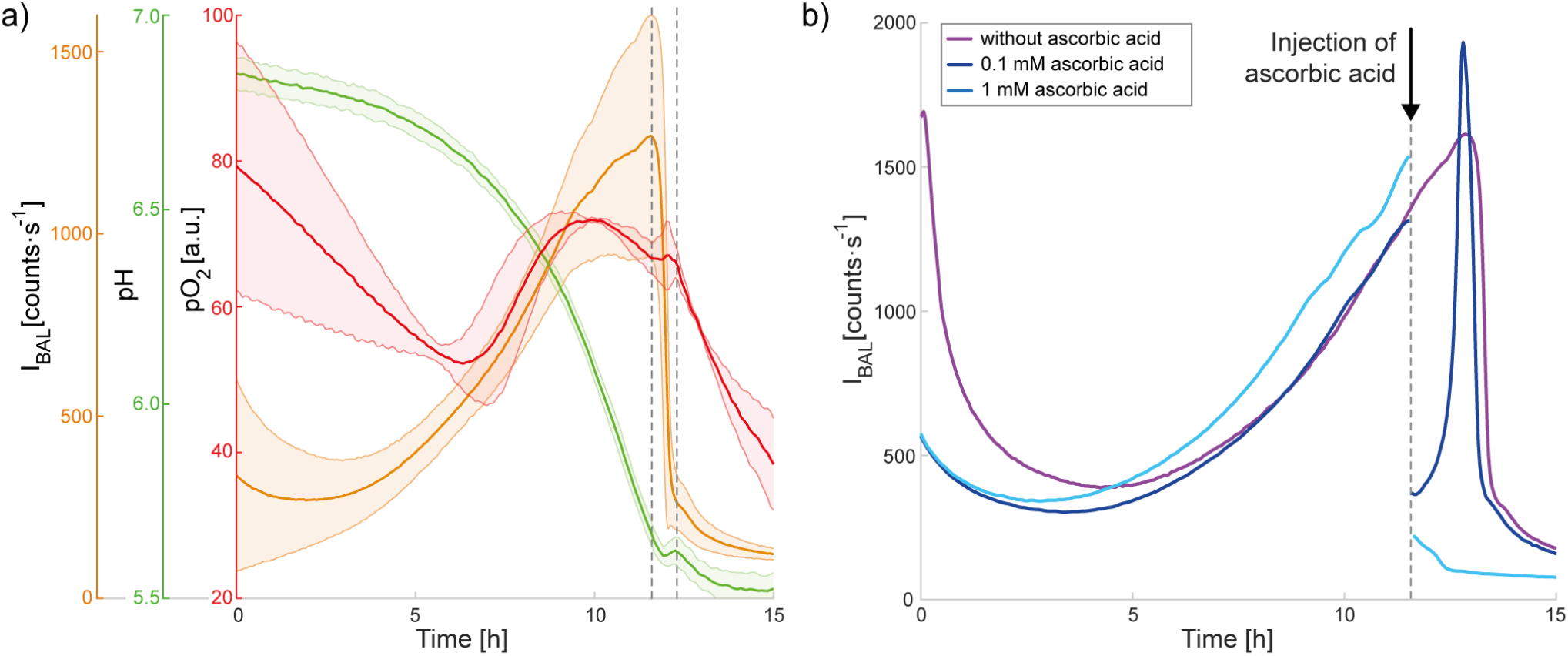
**a)** Spontaneous BAL of yeast measured in a bioreactor together with the change in pH and pO_2_ (average from 3 measurements with a standard deviation). Dashed vertical lines around the twelfth hour delimit the sharp decrease of the BAL intensity and corresponding changes in pH and pO_2_. **b)** Influence of various concentrations of the antioxidant ascorbic acid (0 - 1 mM) on spontaneous BAL of yeast. The initial concentration of the sample was 5·10^5^ cells·mL^−1^.

1. the ROS generation rate 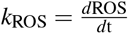 per unit of effective amount of metabolically active yeast biomass *M*_active_ during anaerobic phosphorylation is smaller than the one during the oxidative phosphorylation, i.e., *k*_ROS,anaerobic_ *< k*_ROS,aerobic_
2. the metabolic activity of yeast is much higher during anaerobic metabolism than during aerobic metabolism, as demonstrated by the cell growth-rate in the exponential phase versus stationary phase, i.e., *M*_active,anaerobic_ *» M*_active,aerobic_
3. the BAL intensity *I*_BAL_ is proportional to *M*_active_*k*_ROS_
4. *M*_active,anaerobic_ *k*_ROS,anaerobic_ *> M*_active,aerobic_ *k*_ROS,aerobic_, i.e., *I*_BAL,anaerobic_ *> I*_BAL,aerobic_

What could be the potential sources of ROS during anaerobic metabolism and is there any supporting data for our hypothesis? According to^31^, non-mitochondrial ROS generation can be related to cellular biosynthesis. Tilbury et al. also suggested that possible source of BAL during the exponential phase of yeast growth (during fermentation) are oxidative side-reactions accompanying protein synthesis^11^. Rinnerthaler et al. found that an important non-mitochondrial source of ROS in yeast is a NADPH-oxidase ortholog Yno1p/Aim14p^6^. Maslanka et al. also evaluated the content of pyridine nucleotide cofactors and the ratios of their reduced and oxidised form. They observed a higher ratio of NADPH/NADP^+^ for yeast cultivated in the media supporting aerobic respiration^31^. Besides other functions, NADPH is considered to directly operate as an antioxidant to scavenge free radicals and reduce oxidised biomolecules^32^. Jamieson et al.^33^ observed significantly higher (oxidative) stress resistance during respiratory growth and stationary phase than during fermentation. Because incomplete reduction of O_2_ during respiration is usually considered as the major source of ROS^5^, organisms have developed active, antioxidant-based defense mechanisms, which lead to a decreased amount of ROS in cells or repairing of the damage caused by them. Maslanka et al. observed a dependence of ROS amount and also the ROS source on the type of carbon source/concentration of glucose and on the type of metabolism of yeast *Saccharomyces cerevisiae* wild-type and two mutant strains: Δsod1 and Δsod2^31^. Their results showed that non-mitochondrial ROS sources are an important pool of ROS in yeast, especially during fermentation. They also observed that more ROS are generated by yeast in the medium with higher concentration of glucose when they estimated the level of ROS with dihydroethidium (DHET). Because glucose is an important source of energy for yeast and its concentration in a sample has influence on yeast metabolism, we also performed experiments with glucose. We observed changes in the glucose concentration in the yeast samples during their growth (Fig. S4) and an increase in the yeast BAL intensity when glucose was re-supplied into the sample (Fig. S3). Our data are in agreement with the results of Maslanka et al^31^. The glucose concentration decreased with increasing yeast concentration. In our second mentioned type of the experiment with glucose, after the BAL signal reaches the peak and drops to a low level, the addition of glucose causes a temporary increase of the BAL intensity (Fig. S3).

To briefly comment on the kinetics of the pH and partial pressure of oxygen (pO_2_) in the sample during 15-hour yeast growth in the bioreactor (Fig. 3a): The decrease of pH during fermentation is a well-known effect in yeast biotechnology caused by excretion of organic acids and absorption of basic amino acids^34^. The kinetics of pO_2_ is much more enigmatic: Initially, there is a decrease of pO_2_, then an increase starting after approximately 6-7 hours and then a decrease again after 10 hours. In general, there are two competing processes that determine the pO_2_: O_2_ (air) supply to the bioreactor via an injecting tube through a bubbling output and consumption of the O_2_ in the sample, which explain the three observed phases of the pO_2_ kinetics. The initial decrease of pO_2_ might be due to slowly fading auto-oxidation taking place in the YPD medium, which binds O_2_. The following increase might be due to O_2_ supply starting to dominate over the O_2_ consumption. The decrease that starts after 10 h is likely owing to the onset of cellular respiration.

We carried out further experiments to test whether oxidative reactions are indeed involved in spontaneous BAL generation. The amount of ROS in a sample can be increased by the Fenton reaction, as demonstrated in previous studies^23–25^. On the other hand, the sample ROS content can also be artificially decreased by using, for example, an antioxidant. For experiments in which we wanted to suppress ROS (with an expected consequence of suppressing the BAL intensity), a sample with a higher BAL intensity is more suitable for observing a significant signal decrease. Therefore, the influence of the antioxidant ascorbic acid on the BAL of yeast culture was observed at the time points with high BAL intensity (Fig. 3b. The effect of the antioxidant depended on its final concentration in the sample. After injection of ascorbic acid at a lower concentration (0.1 mM), we observed a rapid decrease followed by a steep increase back to the values expected for a sample without antioxidant treatment. The injection of ascorbic acid at a higher final concentration (1 mM) resulted in a steep decrease and no renewal of the signal. The observed effects probably depended on the ratio of the amounts of ROS and antioxidant. At a lower final concentrations of antioxidant, the molecules of ascorbic acid react with ROS and are quickly depleted. Then, the amount of ROS, and thus the BAL signal, increases again without any hindrance. A higher concentration of ascorbic acid either terminates the chain production of ROS or considerably reduces the amount of ROS for such a long time that the yeast can pass through the fermentation to the aerobic respiration phase. Č ervinková et al.^35^ also studied the effects of ascorbic acid (vitamin C), among other antioxidants, on yeast BAL intensity. They tested concentrations of 1, 5, and 10 mM and also did not observe any renewal of the BAL signal. They also tested the influence of ascorbic acid (1, 500, and 1000 *µ*M) on the BAL intensity of HL-60 cells. Even at the lowest used concentration (1 *µ*M), the signal did not recover. As expected, each sample has a different threshold antioxidant concentration and also a different response to various antioxidants^35^.

## Conclusion

We investigated the correlation between selected physical, chemical, and biological factors and the BAL intensity of yeast samples. The oxidation of a yeast sample using the Fenton reaction caused a higher BAL signal with increasing concentrations of the Fenton reagents. A higher BAL signal was also observed with increasing yeast concentration up to a certain limit, after which the BAL signal decreased again. On the other hand, the addition of the antioxidant ascorbic acid to the sample decreased the BAL intensity for some time depending on the antioxidant dose. We did not observe any damaged yeast cells under a microscope after short-time oxidation, just the tendency of the cells to cluster. However, a higher H_2_O_2_ concentration caused prolongation of the lag phase of the yeast growth.

During yeast culture growth in a bioreactor, the BAL intensity increased together with the number of cells, the pH and glucose concentration decreased, whereas the change in pO_2_ was nonlinear. After approximately 12 hours, when the glucose concentration was low and the yeast samples were undergoing the growth phase change (diauxic shift), the BAL signal sharply decreased.

We demonstrated that the BAL intensity of the yeast sample is proportional to the amount of ROS in the sample. Our data indicated that non-mitochondrial sources of ROS also play an important role in the production of ROS and BAL. We were able to monitor various processes closely connected with the change of ROS content in the studied object by observing changes of the BAL intensity over time. However, the ROS content can be affected by many various factors. Beyond chemical factors, it could be also physical factors such as magnetic^36^ and electric field, particularly pulsed electric field, which is a part of our ongoing work. In general, knowing which other factors can influence the BAL intensity and the mechanisms involved will allow the development of a new reliable diagnostic method.

## Methods

Yeast *Saccharomyces cerevisiae* wild type (BY 4741) was used as a model organism. They were cultivated for approximately 24 hours in an orbital shaking incubator (Yihder Co., Ltd; LM-420D) at 30 ^*°*^C, 180 rpm, in standard YPD medium (1 % yeast extract, 2 % peptone, 2 % D-glucose). Then, they were centrifuged (Heraeus Biofuge Stratos) 2 times and the YPD medium was changed to Q H_2_O. Measurement of yeast concentration was performed with a cell counter (Beckman Coulter, Z2 series) in the range 3 - 9 *µ*m. Yeast *Saccharomyces cerevisiae* has a diameter between 1 and 10 *µ*m. Their mean cell size depends on various factors such as growth temperature^37^, strain, and age^38^.

### Biological autoluminescence measurement setup

#### Induced BAL

Induced emission was measured from a 3 mL of yeast sample in Milli-Q water in a Petri dish (Thermo Scientific, diameter 35 mm). The concentration of yeast was between 10^7^ and 10^9^ cells· mL^−1^. The amount of ROS in the sample was regulated by the Fenton reaction

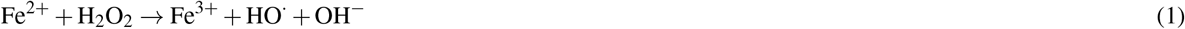

After preparation of a yeast sample at a required cell concentration, 50 *µ*L of iron(II) sulfate was added to the sample so that the final FeSO_4_ concentrations in the sample was 50 nM, 5 *µ*M, or 0.5 mM. Then the sample was placed into a special black box with a photomultiplier module H7360-01 (Hamamatsu Photonics K.K., spectral range 300 – 650 nm) where the measurements of BAL were performed. After approximately 1 min of BAL measurement, 50 *µ*L of H_2_O_2_ (final concentrations: 25 *µ*M, 250 *µ*M, 2.5 mM, and 25 mM) was injected into the yeast sample with FeSO_4_. The whole BAL measurement takes approximately 10 minutes. A scheme of the experimental setup is shown in Fig. 4a.

**Figure 4.**
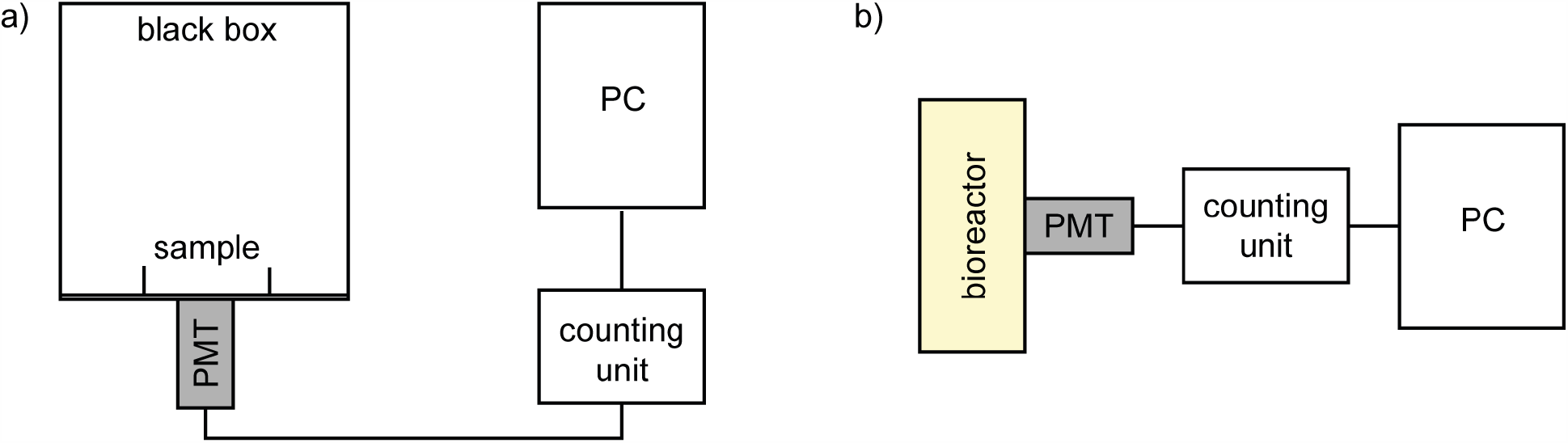
A scheme of the experimental setup for the measurements of BAL during a) induced oxidation in a Petri dish in a black box and b) spontaneous oxidation in a bioreactor.The bioreactor with the photodetector chamber was light-isolated by a black cloth construction.

#### Spontaneous BAL

Spontaneous emission was measured in a bioreactor Biostat Aplus (Sartorius Stedim Biotech) in a dark room. The bioreactor with the photodetector chamber was light-isolated by a black cloth construction to further minimize any background light. Yeasts at an initial concentration of 5 ·10^5^ cells· mL^−1^ were stirred (180 rpm), bubbled with air (∼1 l· min^−1^) and maintained at a temperature of 30 ^°^C during their cultivation in YPD medium. 75 *µ*L of antifoaming agent (polypropylene glycol) was added to the 750 mL sample. The measurement time was between 15 and 20 hours. The photomultiplier module (H7360-01, Hamamatsu Photonics K.K., spectral range 300 – 650 nm) was also used for BAL measurements. A scheme of the experimental setup is shown in Fig. 4b. Measurement of the pH, pO_2_, and temperature was performed by original probes included in the bioreactor.

The initial yeast concentration is lower compared with measurements of BAL intensity after short-time oxidation of a yeast sample in water induced by the Fenton reaction. If we want to measure spontaneous BAL of yeast and their growth curve simultaneously, it is better to start at a lower yeast concentration to have a nicer shape of the curve. (Higher initial amount of yeast depleted glucose quickly, exponential phase is really short, and the difference between the initial and the final cell concentration is not so distinct.) Also, the measurement of yeast concentration is easier and more precise (without dilution of a sample) in the used concentration range. (Upper limit of a sample concentration is approximately 3 ·10^8^ cells ·mL^−1^ for the cell counter.)

The experiment for observing the influence of glucose re-supply into the sample on BAL intensity was performed in an Erlenmeyer flask in a black box similar to the measurements of induced BAL. The measurement setup is described in^39^ and shown in Fig. 1b. Cells were cultivated in an orbital shaking incubator (YPD, 30 ^°^C, 180 rpm) for 16 hours. The concentration of the sample was established with a Bürker chamber. The required amount of yeast cells was transported to 200 mL of cold YPD medium so that the initial yeast concentration was 5 ·10^5^ cells ·mL^−1^. The sample was bubbled with filtered room air. After 16 hours of spontaneous BAL measurement, 11 mL 40% glucose was added into the sample.

### Glucose concentration measurement

The glucose concentration was established with a commercial glucose kit (Glu 1000, Erba Lachema). Optical density was measured at 500 nm and subsequently recalculated to a real glucose concentration in mM (using calibration solutions at known glucose concentration).

### Growth curve

Growth curves were measured in a Spark microplate reader (Tecan). After approximately 24-hour cultivation of yeast in an orbital shaking incubator (30 ^°^C, 180 rpm) in standard YPD medium, the samples ware centrifuged (2x) and the YPD medium was changed to Q H_2_O. Then, oxidation of the yeast sample (concentration of 10^8^ cells· mL^−1^) was performed with the Fenton reaction (0.5 mM FeSO_4_, 2.5 mM / 25 mM / 250 mM H_2_O_2_) for 10 min. The samples were in the shaker (30 ^°^C, 180 rpm) during the oxidation. Then, the samples were centrifuged and the water was changed to fresh YPD medium. After dilution of each sample 100 times, 200 *µ*L of each sample was put in a well of a 96-well plate (Thermo Scientific Nunc, transparent, non-treated, flat bottom, type no. 269620) and covered with a lid. 5-7 replicates of each sample were prepared on the same plate. The optical density at 600 nm was measured every 10 min for 24 hours. The samples were shaken for the rest of the time. The temperature during measurement of the growth curves was maintained at 30 ^°^C.

### Microscopic imaging

Microscopic imaging before and after the oxidation by the Fenton reaction was performed with the microscope BX50 (Olympus) with the camera Moticam 1080. Phase contrast and an objective magnification of 20x was used.

## Supplementary Information

Supplementary Information 1

Supplementary Information 2

## Acknowledgements

We acknowledge Czech Science Foundation, project no. 18-23597S for funding. Authors also participate in the COST CA15211 and exchange project between Czech and Slovak Academy of Sciences, no. SAV-18-11.

## Data availability statement

Raw data are available in the supplementary information 2.

## Author contributions statement

Contribution roles according to CRediT:

https://casrai.org/credit/:

P.V.: Data curation (lead), Formal analysis (lead), Investigation (lead), Methodology (lead), Software (equal), Visualization (lead), Writing – original draft (lead)

K.C.: Data curation (supporting), Formal analysis (supporting), Investigation (supporting), Methodology (supporting)

M.C.: Conceptualization, Formal analysis (supporting), Funding acquisition, Investigation (supporting), Project administration, Resources, Supervision, Validation, Visualization (supporting), Writing – original draft (supporting), Writing – review & editing

