## Supplementary information for "Biological autoluminescence for assessing oxidative processes in yeast cell cultures"

### ABSTRACT

In this supplementary information, we provide additional data to BAL measurements: the autoluminescence signal from media without any cells, data showing the effect of a glucose injection on the BAL intensity, and simultaneous detection of cell and glucose concentration .

#### Water background of (chemi)autoluminescence induced by the Fenton reaction

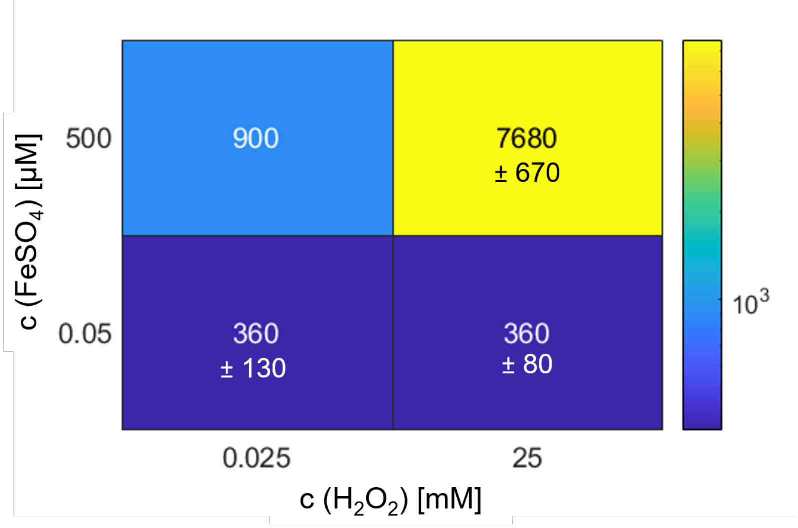


**Figure S1.** BAL from medium alone (Q water without any yeast cells) induced by the Fenton reagents at edge concentrations. The colour indicates the sum of the BAL intensities during the first 60 s after injection of $H_2O_2$ with a subtracted background (the previous 10 s before the injection of $H_2O_2$). The given numbers are usually the average from 3 measurements and the standard deviation.

#### BAL of YPD measured in bioreactor

#### Luminescence recovery after addition of glucose

In Fig. S3 the effect of addition of glucose into the yeast sample after sharp decrease of the BAL signal (approximately at seventeenth hour of the measurement) is shown. The measurement protocol was little bit different than for other experiments. Yeast cells were cultivated in an orbital shaking incubator just for 16 hours (standart YPD, 30 °C, 180 rpm). Yeast concentration was established using Bürker chamber. Required amount of cultivated yeast was transferred to 200 mL of cold YPD medium at initial concentration $5{\cdot}10^6$ cells$\cdot$mL$^{-1}$ placed in Erlemeyer flask covered with cotton stopper. The sample was bubbled with filtered room air. The experimental setup for BAL measurements was similar to the setup for measurements of induced

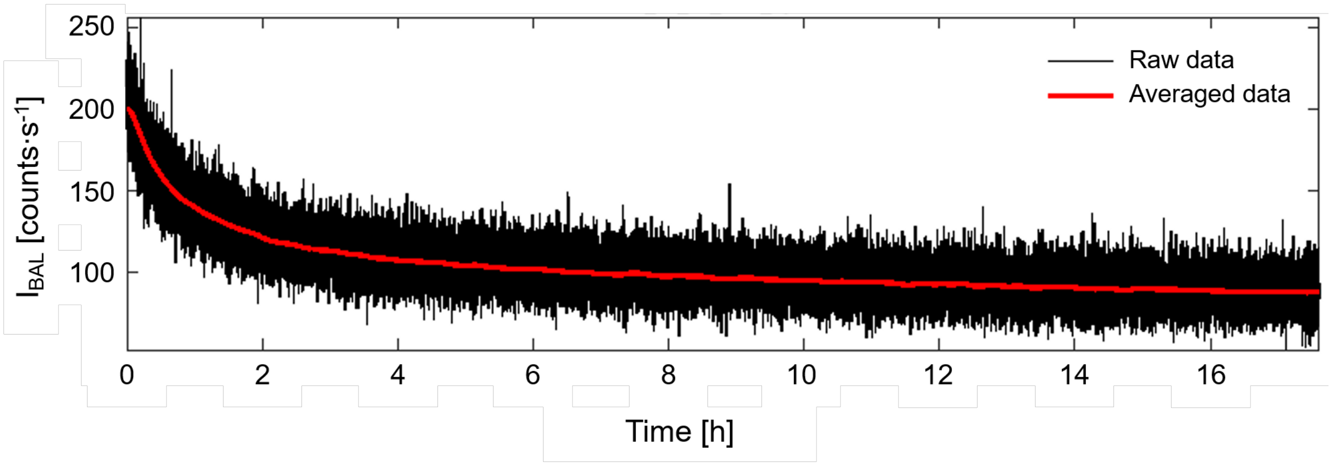


**Figure S2.** BAL signal from YPD medium (without yeast cells) measured in bioreactor for almost 18 hours. Initial higher BAL intensity was caused by the illumination of the sample during its preparation in the normal laboratory light conditions.

BAL from samples in a Petri dish, just instead of a Petri dish an Erlenmeyer flask was placed in a special black box (30 °C) with a photomultiplier module H7360-01. After the sharp decrease of the BAL intensity (approximately after 17 hours of measurement), 11 mL of 40% glucose was injected into the sample. The measurement was several times interrupted in order to establish the yeast concentration.

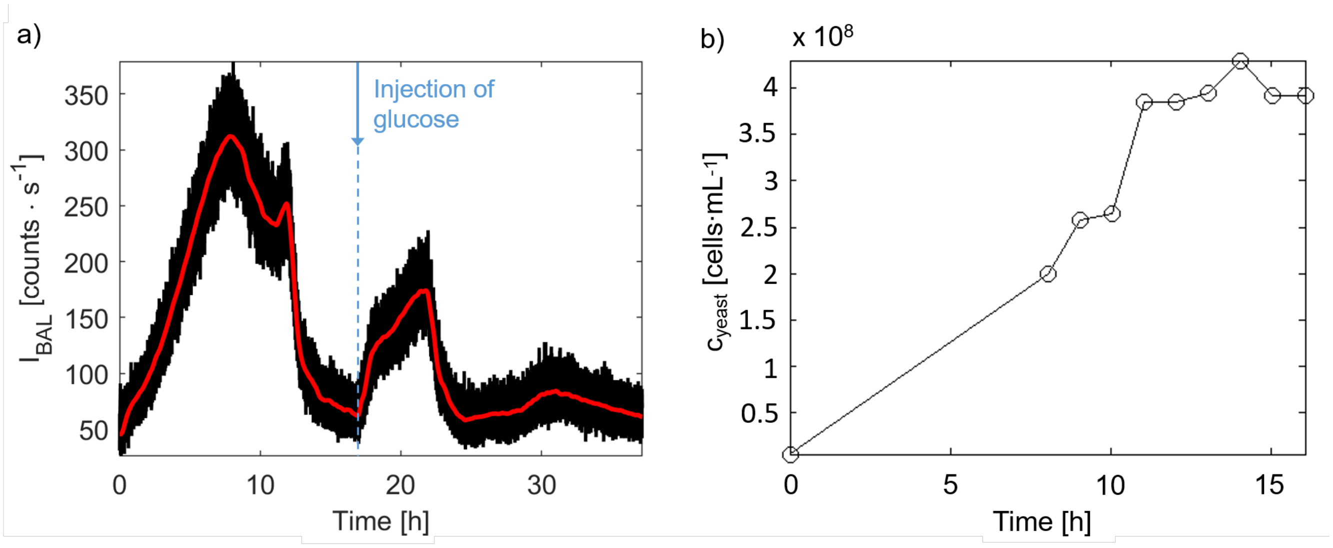


**Figure S3.** Development of BAL intensity during yeast growth in Erlenmeyer flask and repeated increase of the BAL signal after injection of glucose.

### Development of glucose concentration in a growing yeast culture

Fig. S4 shows the development of glucose concentration during the yeast culture growth in bioreactor that is depicted by yeast cell concentration. Yeast concentration was evaluated by a cell counter (Beckman Coulter, Z2 series) in the range 3 - 9 $\mu$m. Glucose concentration was established by a commercial glucose kit (Glu 1000, Erba Lachema). Optical density was measured at 500 nm and subsequently recalculated to a real glucose concentration in mM (using calibration solutions at known glucose concentration). Both measurements are subject to a large deviation. However, trends are clear: with increasing amount of cells in the sample, glucose decreases. In other words, low concentration of glucose is correlated with the change of yeast growth phase and the change of yeast metabolism.

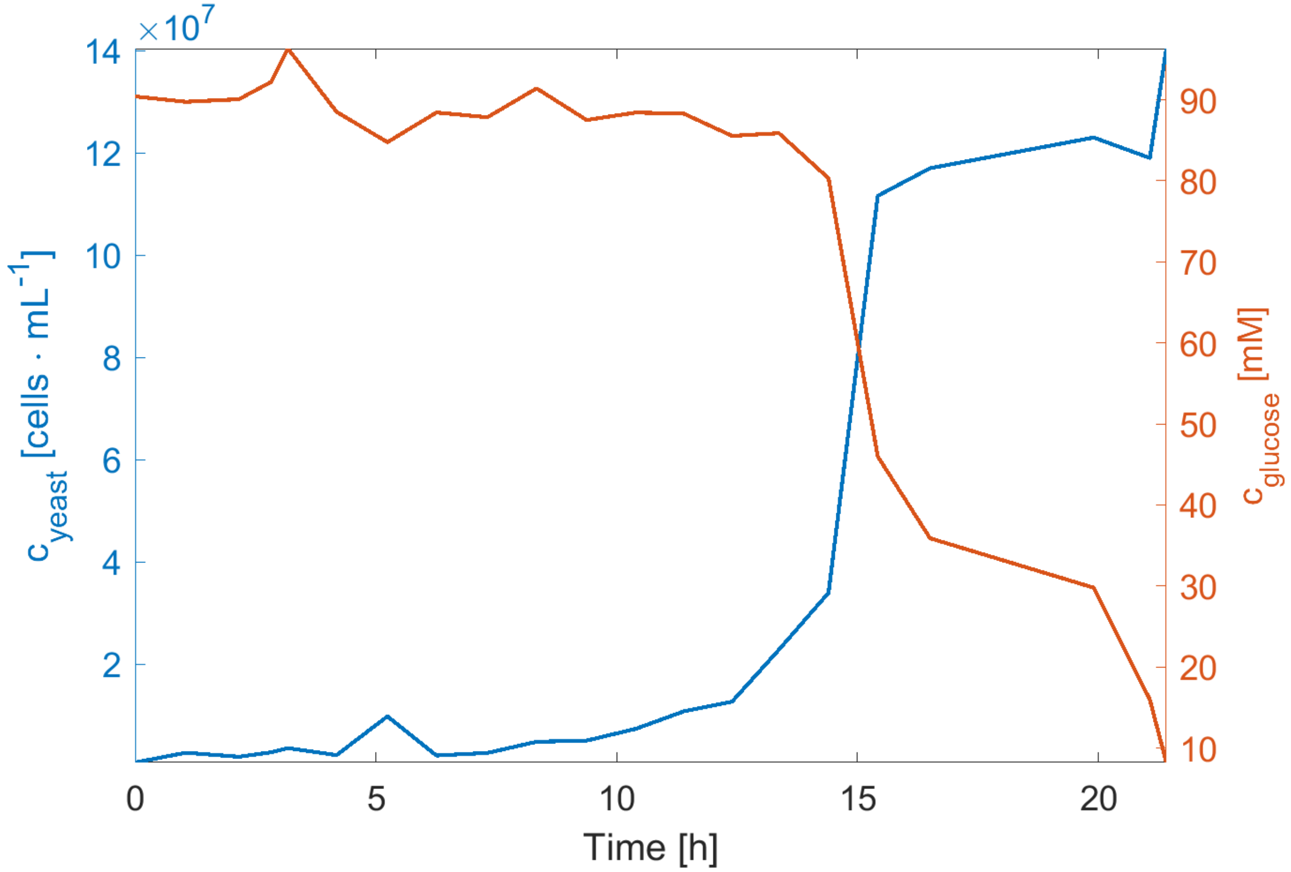


**Figure S4.** Simultaneous measurement of concentrations of glucose and yeast cells during their growth in a bioreactor. While the small measurement errors are visible, the trend is clear.

### Raw data description

The source data can be found in the Supplementary information 2 ( .zip file) and are sorted according to the figure number and part. For plotting the mean and the standard deviation of several (3 - 7) independent measurements in the Fig. 3a, 3d, and 4a, the predefined Matlab function 'shadedErrorBar' was used. Figures 4(ab), S2, and S3 contain smoothed data obtained by function 'rsmooth'[1]. To create Fig. 3c and S1 the Matlab command 'heatmap' was used.
